# Classification of Communication and Head Movement Behaviors during Multi-Person Conversations using Deep Learning

**DOI:** 10.1101/2025.09.26.678869

**Authors:** A Earley, A Chhabra, EJ Ozmeral, A Lertpoompunya, DA Eddins, NC Higgins

## Abstract

Head movements play a pivotal role while engaged in multi-talker conversation by providing non-verbal feedback to partners and enhancing a listener’s ability to separate sound sources. Commercial hearing-aids with an on-board IMU (Inertial Measurement Unit, i.e., accelerometers) typically use that information for step-counting and activity levels. At least one device uses it as an input to their environment classifier and integrated directional microphone. None, however, use the IMU to detect specific patterns of head movements to predict behaviors such as nodding, head shaking, listening to a person versus a video, or talking. Training an automatic classifier to accurately detect these behaviors first requires collecting head movement data during multi-person conversation and laboriously annotating each behavior-type for each participant with high temporal precision. From that point, with the goal of training the most accurate model and integrating with the hardware in the hearing aid to improve device performance, the question is how best to model the data. To address this gap, we collected accelerometer data during natural multi-person conversations and paired it with detailed human annotations of communication and head-movement behaviors. Head-movement data was collected from three cohorts of young, normal-hearing individuals (three per cohort) in a controlled, conference-room setting during 50-minute multi-talker conversations. Participants wore hearing aids with on-board accelerometers, and audio-video was recorded for each talker. Videos were manually annotated for communication and head-movement behaviors, including conversational turns and nonverbal cues such as tilts and nods. Temporal and spectral features were extracted from the accelerometer data (windowed into 1-second segments) corresponding to roll and pitch movements. These features, combined with the annotated behaviors, were used to train and test machine learning models. Models were trained on data from all but one participant and then tested on the held-out participant, repeating this procedure across all individuals. Several deep learning and classical machine learning models were compared for classifying communication behaviors (e.g., talking, listening, watching video) and head orientations (e.g., turning left or right, facing down, facing forward). More specifically, various sequence-to-sequence models, a state-of-the-art deep learning technique, were utilized. These models incorporated modern architectural components such as transformer networks. Multiple performance metrics were used to evaluate models, and results suggest that modern deep learning models outperform classical machine learning methods by significant margins. Classification performance improved further when temporal sequence information was incorporated. These results indicate that during multi-talker conversation, hearing-aid accelerometers can automatically classify stereotypical behaviors with high temporal resolution (1-second). Even when tested on unseen subjects, the models remained reliable. This establishes a foundation for more advanced approaches that combine behavioral and movement patterns, further integrate temporal dynamics, and incorporate additional inputs to improve accuracy and ecological validity.

## Introduction

Advanced hearing aid technology includes various sound cleaning technologies designed to improve the listening and communication experience. Among those, spatial hearing systems (directional microphones or microphone arrays) are mainly used to improve the signal-to-noise ratio (SNR) for the listener, with the goal of better speech intelligibility in noisy environments. However, benefits associated with those technologies come with the cost of reducing binaural acoustic cues, resulting in a tradeoff between SNR improvement and binaural cue preservation (J. G. Desloge et al., 1997; Van den Bogaert et al., 2006; Doclo et al., 2010; Ibrahim et al., 2013; Neher et al., 2017). Importantly, however, some spatial hearing systems can locate the spatial position of a talker and selectively amplify sounds from that location more than sounds from other locations (i.e., beamforming). In realistic communication situations, however, this strategy is prone to error when listeners move their heads and the beamformer fails to maintain the target stream or adapt to the new location. Accurate tracking of target talkers is particularly problematic in multi-talker environments when a listener is likely to move their head away from the talker or switch from talker to talker at the same rate as the conversation. Despite sophisticated real-time signal processing in modern premium hearing aids, current devices are unable to recognize user intention (e.g., actively listening versus talking) and cannot adapt to communication behaviors such as head and eye movements.

Natural listening situations often involve multiple sources, and those source locations are not always constant over time. As a result, hearing aid users may have difficulty 1) developing awareness of sound source locations, 2) ignoring irrelevant sounds, 3) changing focus from one source to another, and 4) tracking moving sources. Head orientation and movement patterns are a prime candidate to source the information needed to identify listening intentions and ultimately, to improve hearing aid algorithm performance (Kidd et al., 2013, 2015; Best et al., 2017; Favre-Felix et al., 2017; Kidd, 2017; Favre-Félix et al., 2018; Roverud et al., 2018; Hladek and Seeber, 2019; Lu et al., 2021). Conveniently, newer hearing aids often have on-board accelerometers capable of capturing head movements, opening the door to the integration of head movement information to the hearing aid processor to adjust the sound treatment strategy in real-time. Development of a “smart” feedback system would first apply principles in theories of social interaction, auditory scene analysis, and listening behaviors to predict who, what, and where the device should be enhancing, as indicated by a user’s listening intent.

To systematically examine human behaviors Wolters et al. (2016) proposed the common sound scenarios (CoSS) framework for evaluating three categories of listener intent: 1) speech communication, 2) focused listening, and 3) non-specific hearing. In addition, several studies have examined listening situations that hearing aid users encounter in daily life (Wagener et al., 2008; Wu and Bentler, 2012; Wolters et al., 2016). Using the CoSS framework, (Smeds et al., 2015) reported that almost half of the time, hearing-impaired listeners categorized their listening intent as *nonspecific*, whereas only a third of the time was categorized as speech communication and 20% was categorized as focused listening absent of active speech communication (e.g., television or radio). Hearing aids, however, may treat these situations quite similarly, and considering these proportional differences in listening intent, perhaps it is not surprising that many hearing aid users are unhappy with their aided listening (Appleton, J., 2022; Desai et al., 2024).

One of the essential components of developing an experiment with realistic communication is participation or engagement in the conversation itself, meaning multiple participants are needed to fully engage behavioral systems that not only promote audio and visual perception but also maintain certain social norms. To that end, we devised an experimental scenario that included multiple conversational partners and prompted them to engage in lively discourse. During the discussion we recorded multiple aspects of the environment, including audio, video, head movements, and eye gaze. Based on the audio-video recordings, the CoSS framework was then used to annotate (label) specific head movements and communication behaviors, and time-match them to physio-kinetic measurements related to head movements recorded from hearing aid-based accelerometers.

The resulting dataset can then be used to train an automatic classifier to determine stereotypic head movement behaviors that correspond to elements of listening intent. With the rapid introduction of machine and deep learning algorithms, the immediate goal of this report is to determine and compare different algorithms and identify the best-performing, with the end-goal of porting the model to a hearing aid simulator.

In general terms, artificial intelligence is the idea that machines can be made to imitate human learning and reasoning (Russell et al., 2020). Machine learning specifically, is a branch of artificial intelligence in which models improve automatically by learning from data. This is important because many tasks are too complex to be hand-coded, and learning from data allows systems to adapt and generalize to new situations (Russell et al., 2020).

Deep learning is a subset of machine learning that excels at working with larger amounts of data, training them over a series of steps (Russell et al., 2020). During this time, the model aims to minimize the prediction error, known as loss. These models use their validation sets to provide periodic checks and tuning during training, with one full pass through the training set referred to as an epoch (Russell et al., 2020).

As such, both classical machine learning and deep learning techniques provide powerful tools to analyze large amounts of data and extract meaningful patterns very quickly. Many models provide even more specialized tasks that are also applicable to biological systems. For example, Recurrent Neural Networks (RNNs) and sequence-to-sequence models like the Mambular model are deep learning models that are especially good at predicting time-dependent data (Elman, 1990; Thielmann et al., 2024), ideal for patterns of behavioral activity.

Altogether, this paper aims to address the hypothesis of whether various classical machine and deep learning models can accurately predict head movement and communication behaviors from hearing aid accelerometer data. In addition, it aims to compare their relative performance to determine the future design of real-time wearable devices able to accurately predict listener intent.

## Methods

### Participants

A total of nine young normal-hearing participants, ranging in age from 20 to 35 years old (mean 25.3; standard deviation 5.2), participated in this study. Participants were tested in cohorts of three individuals (3-people per group x 3 groups) who did not know each other prior to participation. Inclusion criterion for normal hearing was defined as pure-tone thresholds less than 25 dB HL at octave frequencies from 250 to 8,000 Hz. Inclusion criteria also included the ability to complete the behavioral tasks at any level of performance and a score on the Montreal Cognitive Assessment (MoCA) test of 26 or above (Nasreddine et al., 2005). This experiment was a one-visit study. All procedures were approved by the University of South Florida Institutional Review Board. Written informed consent was obtained from all participants, and participants were paid for their participation.

### Equipment and Apparatus

To model communication environments characteristic of natural settings, the experiment was conducted in a 3 m x 4 m room (Fig. 1) at University of South Florida. Three participants were included during each experiment session and were situated around a circular table at 120° intervals. Each participant was equipped with the following for physio-kinetic measurements:

**Figure 1.**
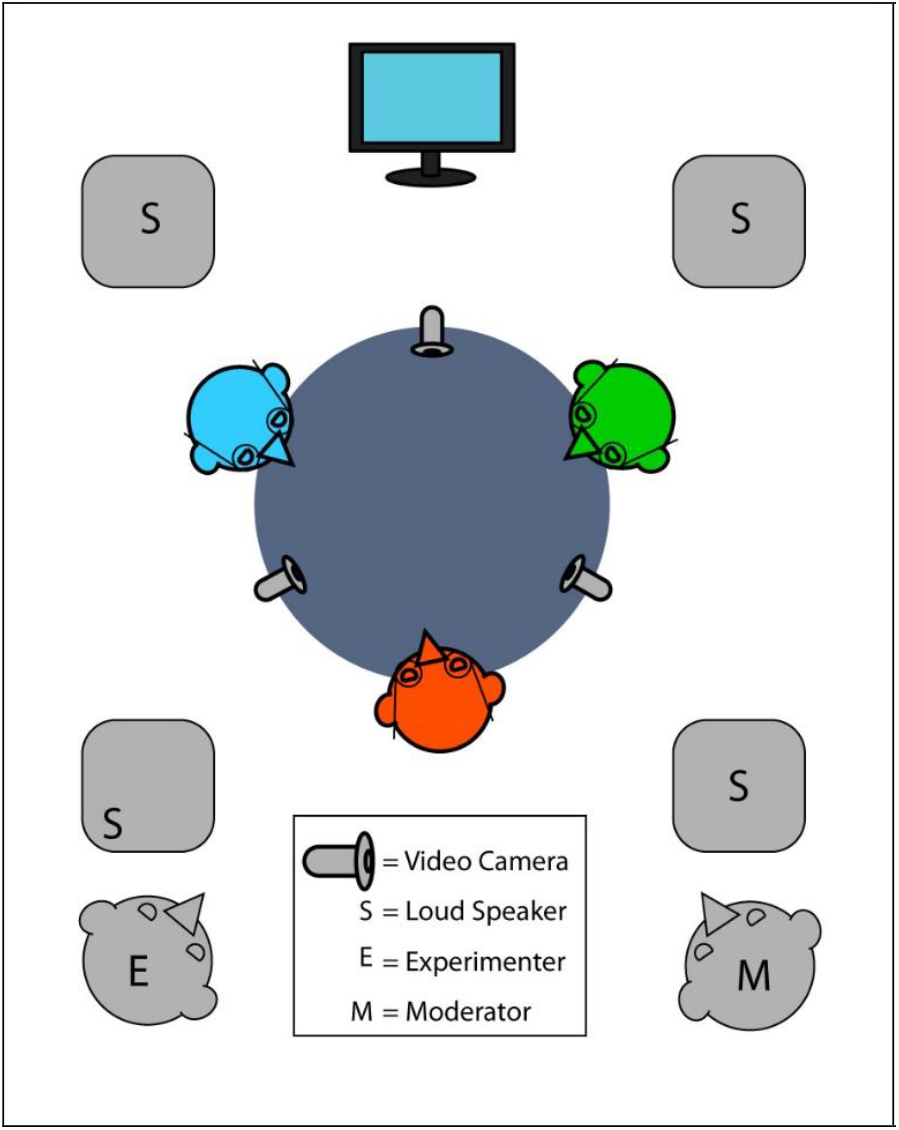
Experiment design. Three participants (labeled red, green, blue) were seated around a circular table. Each wore a head mount with markers for the infrared camera system (OptiTrack V120-Trio, yellow), Pupil Lab eye tracking glasses, and hearing aids equipped with accelerometers. Three video cameras individually recorded each participant. Four loudspeakers (squares with “S”) surrounding the participants delivered background noise during the different experimental conditions. The TV monitor was in front of the room. A moderator (“M”) and an experimenter (“E”) were sitting in corners of the room so that they did not block the camera view. A moderator was available to provide conversational prompts when necessary.

1. Head mounted array with attached infrared sensors; movement of the sensors was tracked with the OptiTrack V120-Trio camera.
2. Pair of hearing aids (Unitron MoxiB90) configured to provide no amplification with an open fit (no occlusion). Each hearing aid (left and right ear) contained an accelerometer.
3. Pupil Invisible glasses that recorded video from the wearers’ perspective and provided eye-tracking throughout the experiment. These devices also provided head movement data.

Across the table from each participant a camera recorded each person throughout the experiment (Reolink 4k). Video for each participant was subsequently used for annotation of behaviors and calculations of yaw, roll, and pitch head movements using z-face software (Jeni et al., 2017). In total, head movement data was captured from four systems: the OptiTrack system, the Pupil Invisible glasses, the video recording, and the hearing aid accelerometers. Post-experiment analysis showed high agreement between all systems. Verification and synchronization between the zface-video data (yaw, pitch, and roll) and accelerometer data were used to verify that the time-axis of the video annotation and the accelerometer measured were aligned. Note that the hearing aid accelerometers only provide roll and pitch head movements.

### Stimuli

An audio-video clip (∼3 minutes) was presented from the TV monitor (Fig. 1) in front of the room for the purpose of generating a common basis for subsequent conversation. Two background noise conditions were tested: 1) broad band noise (BBN) at 60 dB(C) and 2) multilanguage-babble at 60-dB(C), in addition to ambient room noise. Background noise was presented from four loudspeakers (Behringer Truth B2030A) positioned in a square around the participants equidistant from the center of the table, as shown in Figure 1. There were six video sessions in total, three with BBN and three with babble. The order of the background noise condition was counterbalanced between participant groups. A tone was presented every ten minutes to aid in post-processing and synchronization.

### Procedures

To simulate natural conversation, the three participants were seated around a circular table, spaced equally apart (Fig. 1). Participants watched an audio-video clip (∼3 minutes, topics: food, travel, car, hearing aids, and music) presented from a monitor that is visible and audible to each participant for the purpose of generating a common basis for subsequent conversation. A notepad was positioned on the table in front of each participant at the beginning of the session and during the audio-video clip they were allowed and encouraged to take notes as needed to facilitate a subsequent conversation about the audio-video clip. After watching the clip, participants had ∼2 minutes to write down answers to guided questions or any other ideas that they would like to discuss with the other two session participants. Following the note-taking segment, participants were asked to engage in conversation for 5-minute period. They were free to discuss anything regarding the video (prompted by moderator, if necessary, e.g., asking a question or introducing an idea about the video) and to pursue any tangents that arose from that topic. During the conversation, they could freely move their heads. The entire session was audio and video recorded using a separate camera focused on each participant. Those recordings were used for coding purposes to match types of behaviors to the head movement data.

### Behavioral Annotation

The entire session was audio and video recorded and subsequently annotated to match types of behaviors to associated head movement data. Annotation was based on three primary behavioral categories: a) head movement: facing down, facing forward, nodding, shaking, turning to right, turning to left, b) aural-oral communication: talking, listening to talker, listening to video; c) others: writing, looking at cell phone, fidgeting. Three annotators coded the recordings independently using the EUDICO Linguistic Annotator (ELAN) 6.4 software application (Lausberg and Sloetjes, 2009). Audio-video recordings from each participant were imported to ELAN and annotated by each annotator individually frame by frame. Each behavior was segmented by onset and offset times in segmentation mode.

In natural conversation, people constantly move while they are talking or listening, and consequently, head movement and aural-oral communication can overlap in time. Therefore, annotations were grouped in tiers based on behavioral category. By rule, two or more behaviors in the same tier (i.e., the same behavioral category) could not overlap, while behaviors in different tiers could overlap. For example, turning the head to the right could be coded as occurring at the same time as talking. During transcription mode, annotators characterized which behavior was observed based on a pre-defined taxonomy of behaviors (Table 1), and the time segments over which those behaviors occurred were marked. The 16 possible behaviors were defined as controlled vocabularies and linked to a behavioral category, or tier, then selected from pre-defined drop list of labels. Events tagged by the annotators for certain behaviors such as head shaking, nodding, talking, and listening, were time-aligned and co-registered with the head tracking data in post-processing by matching trigger tone. One behavior that was ultimately excluded from the analyses was “looking at cell phone” as it was never observed.

**Table 1.** Pre-defined taxonomy of behaviors.

| **Behavioral category / tier** | **Controlled vocabulary** | **Action** | **Definition** |
| --- | --- | --- | --- |
| Head movement | FaceDown | Facing down | Face pointing downward |
|  | FaceForward | Facing forward | Looking straight ahead, facing to the front |
|  | Nod | Nodding | Repeat sweep on Y axis (pitch) |
|  | Shake | Shaking | Repeat sweep on Z axis (yaw) |
|  | TurnL | Turning to left | Head rotation oriented to the left (yaw) |
|  | TurnR | Turning to right | Head rotation oriented to the right (yaw) |
| Aural-oral communication | TalkL | Talking to the left | Actively talking to participant on the left |
|  | TalkR | Talking to the right | Actively talking to participant on the right |
|  | TalkU | Talking undefined | Actively talking to a mix of listeners |
|  | ListenL | Listening to the left | Active listening to another participant on the left |
|  | ListenR | Listening to the right | Active listening to another participant on the right |
|  | ListenU | Listen undefined | Active listening to a mix of talkers |
|  | ListenV | Listening to video | Active listening to video |
| Others | Note | Taking note | Jot down, writing notes |
|  | Phone | Looking at cell phone | Looking at or using cell phone |
|  | Fidget | Fidgeting | Restless movement of head or body |

### Data Analysis, Head Movement

Roll and pitch head movement trajectories, collected by the accelerometers in the hearing aids, were time-aligned with roll and pitch trajectories from the video-based zface analysis (Fig. 2). Once the head movement signals were synchronized in time, the dataset representing labeled behaviors matched to roll and pitch head movements were segmented into 1-second samples. From each 1-second roll and pitch sample, features were calculated corresponding to: peak velocity, root mean square, standard deviation, mean, linear slope, frequency of sine wave fit, residual of sine wave fit, frequency from FFT (fast-Fourier transform), magnitude of the peak of the FFT, sequential order of the 1-second sample within the labeled behavior.

**Figure 2.**
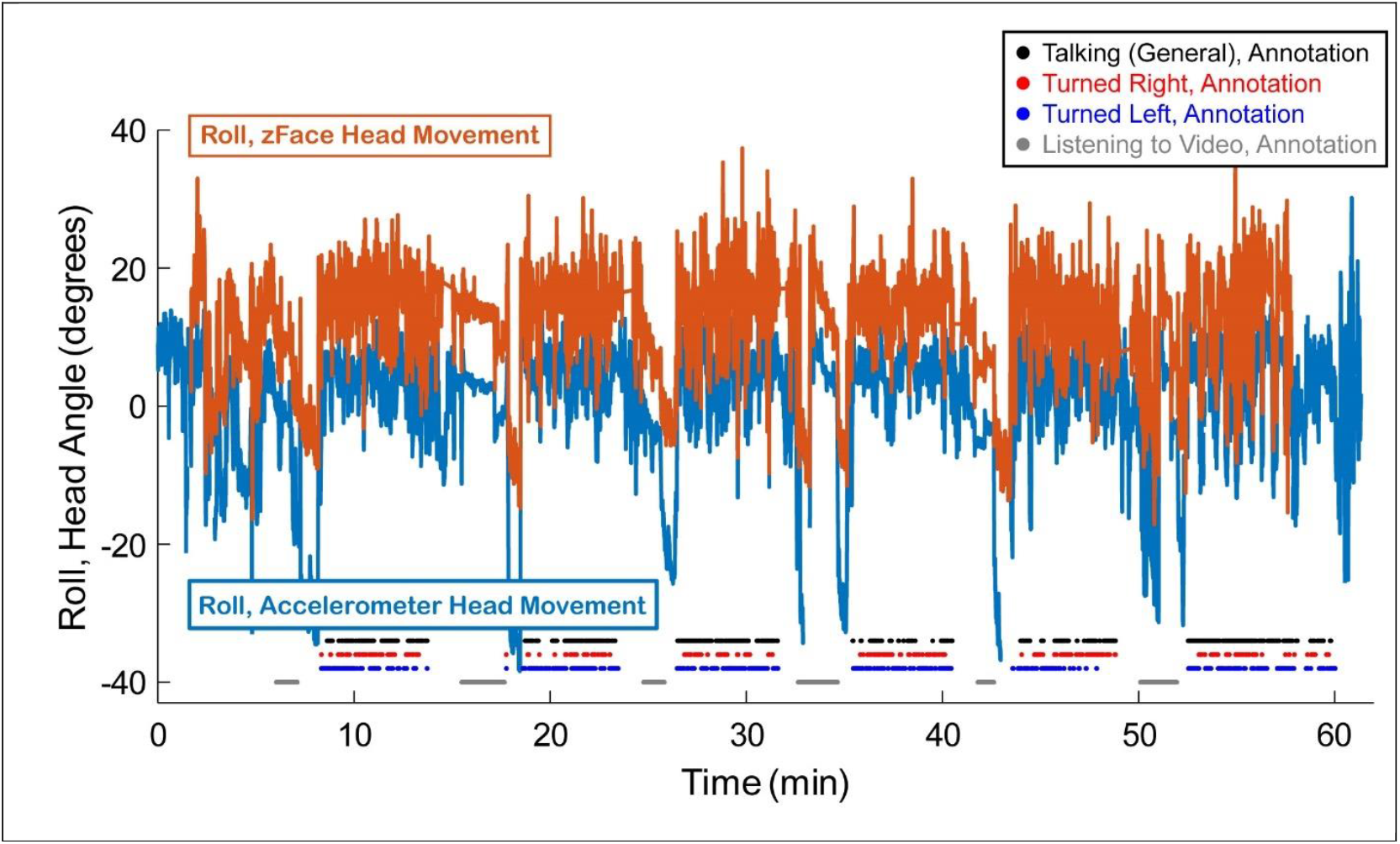
Example data from one participant of time synchronized Roll head movement collected from the zFace video calculation (orange) and the Roll head movement measured by the hearing aid based accelerometer (blue) over the 1-hour experiment. Markers along the bottom of the panel represent labeled behaviors in 1-second segments corresponding to annotations of Talking (black), Turned to the Right (red), Turned to the Left (blue), and Listening to the Video. Note that periods of watching/listening to a video were typically followed by a sharp decrease in the Roll, corresponding to periods where the participant put their head down to write. Discontinuities in the Accelerometer data occurred between video segments when the device was turned off and restarted to assist with time-synchronization.

### Classification Methods

Various classical machine learning and deep learning models were used to predict these communication behavior and head movement features. To create our models, we compared classical machine learning approaches with deep learning models to evaluate which performed best. For a baseline, we included scikit-learn’s Dummy Classifier (Pedregosa et al., 2011), which provides an untrained reference point. A diverse set of models was chosen to avoid biasing the comparison toward any particular approach.

The classical models included:

- Linear Support Vector Machine (SVM) (Cortes and Vapnik, 1995)
- Support Vector Machine with a Radial Basis Function kernel (RBF SVM) (Cortes and Vapnik, 1995)
- Random Forest classifier (Breiman, 2001)

The deep learning models, implemented using the Mambular library (Thielmann et al., 2024), included:

- Multi-layer Perceptron (MLP) (Rosenblatt, 1958)
- ResNet (Residual Network) (He et al., 2016)
- Tabular Recurrent Neural Network (RNN) (Elman, 1990; Hochreiter and Schmidhuber, 1997; Thielmann et al., 2024)
- Mambular (Thielmann et al., 2024)

All models were initialized with their default hyperparameters, with the most notable ones shown below in Tables 1 and 2 for completeness. These values allowed for a solid basis for initial comparison. In addition, the deep learning models used cross-entropy loss. This approach is the standard way to measure prediction error when a model differentiates between multiple classes. Furthermore, all deep learning models used an early stopping patience of 10 epochs. In other words, if the performance stopped improving on a separate set of data held out during training (the validation set, used only for checking progress), training stopped and the best-performing model was restored.

For the classical models, we utilized k-fold cross-validation with *k* = 9 (Kohavi, 1995). Specifically, we applied the Leave-One-Subject-Out (LOSO) strategy, where each subject was tested on models trained using the remaining subjects. The deep learning models followed a similar setup: of the 8 non-testing subjects, 1 was randomly assigned to the validation set, while the other 7 comprised the training set. This strategy ensures testing is robust and not biased towards any particular subject. By testing on an unseen subject, this evaluation procedure makes the results more representative of how the model would perform on new individuals in real-world use, improving ecological validity.

**Table #:**
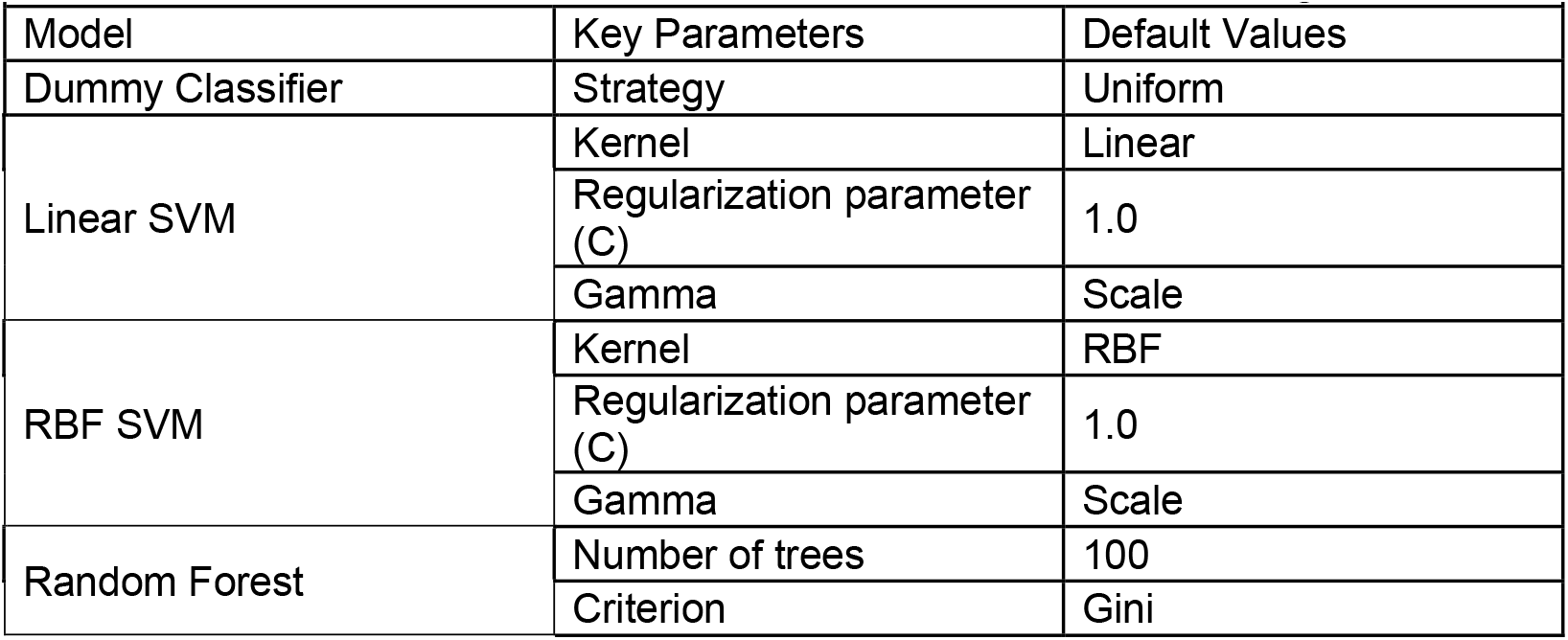
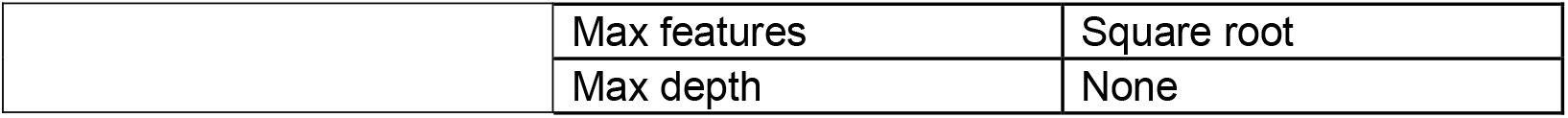
Parameters and Default Values for Classical Machine Learning Models.

**Table #:** Parameters and Default Values for Deep Learning Models.

| Table #: Parameters and Default Values for Deep Learning Models |  |  |  |  |  |  |
| --- | --- | --- | --- | --- | --- | --- |
| Model | Layers / Blocks | Hidden Dimension | Dropout | Activation Function | LR | Weight Decay |
| MLP | 3 layers (256,128,32) | 32 | 0.2 | ReLU | 0.0001 | 1e-6 |
| ResNet | 3 residual blocks | 32 | 0.5 | SELU | 0.0001 | 1e-6 |
| RNN | 4 layers | 128 | 0.2 (RNN), 0.5 (head) | ReLU (main), SELU (head) | 0.0001 | 1e-6 |
| Mambular | 4 layers | 64 | 0.0 (main), 0.5 (head) | SiLU (main), SELU (head) | 0.0001 | 1e-6 |

Note: ReLU = Rectified Linear Unit; SELU = Scaled Exponential Linear Unit

Model performance was evaluated using a variety of F1 scores. These metrics are a more balanced measure than accuracy, which can be skewed by unequal numbers of false positives and false negatives. First, F1 scores were calculated for each class within the behavioral communication or head movement category. From these, we then derived two summary metrics: Macro-F1 and Weighted-F1. The Macro-F1 score is the unweighted average of the class F1 scores, giving equal importance to each one. This is particularly useful when all classes are considered equally important. For example, our dataset contained slightly fewer “watch video” observations than other communication behaviors, yet Macro-F1 gave it equal weight, highlighting performance on minority classes as much as on majority ones. In contrast, the Weighted-F1 score averages class F1 scores according to their relative frequencies, providing a performance measure that reflects the class distribution.

## Results

In total, the data collection procedures resulted in recorded multi-person communication for three triads. Each triad contributed 50 to 60 minutes of raw data. Labeling of the full dataset for head movement and communication behaviors was completed independently by two annotators.

### Annotation

Following full annotation of data from all participants by two annotators, action segments were converted from on/off time stamps to a continuous time-axis at a resolution of 100 Hz. Figure 3 illustrates an example from one participant during the hour-duration experiment and all corresponding labeled action segments from one annotator. Experimental conditions corresponding to separate video periods can be observed from the Listen to Video label when video prompts were presented. Note that during those time periods all other labels were typically *off* except for Face Forward. In between these video segments clear periods of conversation were observed as indicated by the high counts of action segments corresponding to Head Movement and Communication Behaviors. The behavior label for “Phone” was never identified by either annotator for any subject and is not included in further analysis.

**Figure 3.**
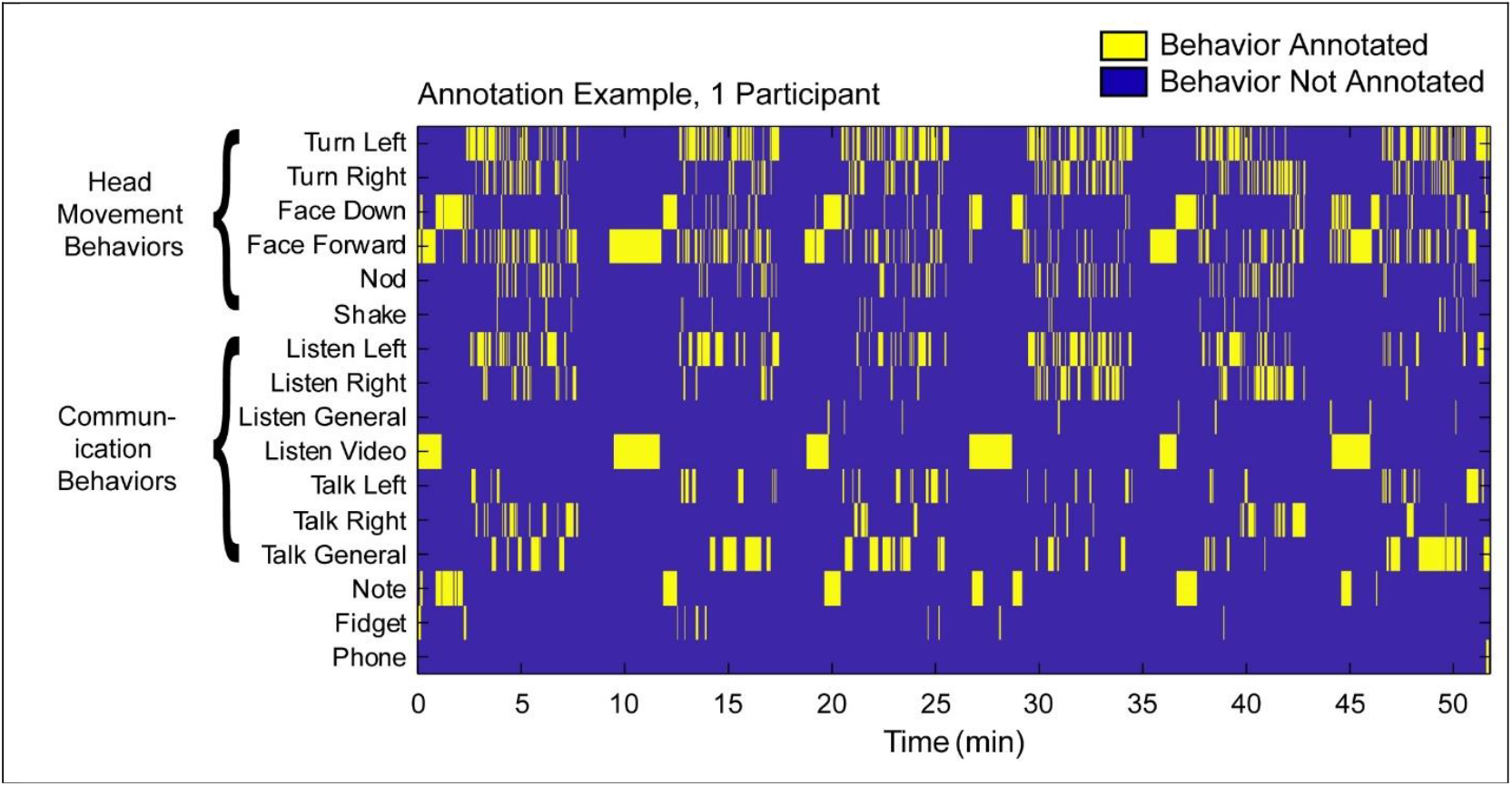
Timeline from one participant of all labeled behaviors as identified by one annotator. Time points representing onset and offset of each segmented behavior were converted to a continuous binary code for each label with annotated behavior in in yellow and periods of time when that behavior was not observed, in blue.

Boxplots representing average frequency counts and average time durations for each behavioral label (Fig. 4) show that Head Movement Behaviors were observed more often than Communication behaviors. Overall, annotator 2 identified more segmented behaviors than annotator one, but the overall pattern between the annotators was very similar (Fig. 4 A and B). and high agreement between the two annotators was observed (Fig. 5).

**Figure 4.**
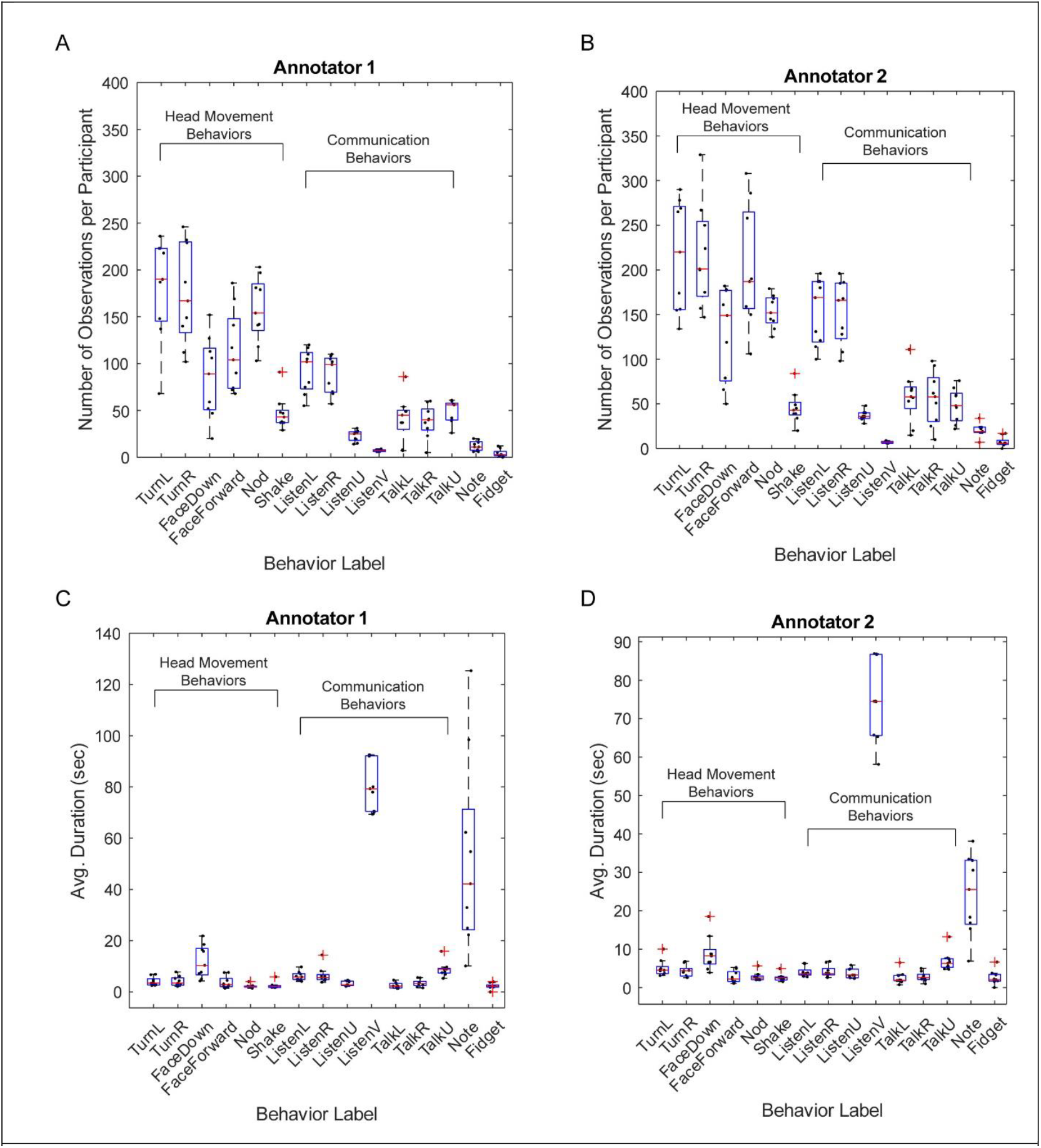
Overview of annotated behavioral labels. A) Frequency count of labeled behaviors by Annotator 1, and B) Annotator 2. C) Average duration of each labeled segments for Annotator 1 and D) Annotator 2. Box plots correspond data measured across the 9 participants: top and bottom edge of each box represent the 25^th^ and 75^th^ percentiles and the red line is the median. Black dots represent data from each participant.

**Figure 5.**
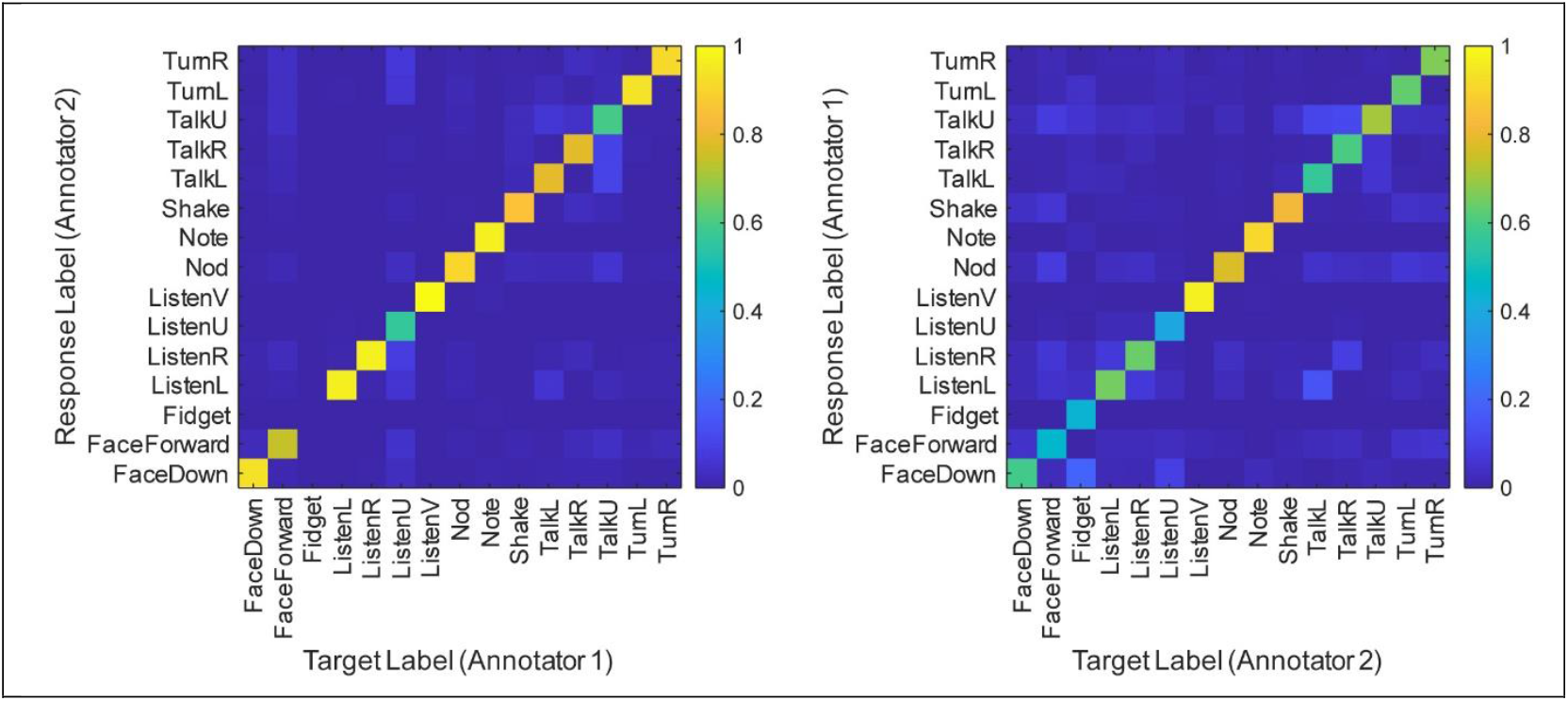
Confusion matrices representing the probability of overlap for each behavioral-label between Annotator 1 onto Annotator 2 (left panel) and Annotator 2 onto Annotator 1 (right panel).

### Classification Results

Following annotation, the input and output features were combined to balance performance and ecological validity. The final model inputs consisted of peak velocity, root mean square, mean, and standard deviation of both roll and pitch, yielding a total of eight features. Sequence length, defined as the duration of the action in seconds, was included as an additional input feature.

The model then classified communication behaviors into four categories: talking, listening, watching a video, or other behavior type. Head movement outputs were grouped into facing forward, turning left, turning right, facing down, or other movement. Models were trained as described in the *Methods* section.

Figure 6A shows mean training loss curves for deep learning models trained on communication behavior data, averaged across nine subjects. Training loss, which measures how far off the model’s predictions are from the correct answers, declined over time, with the steepest drops observed in the RNN and Mambular models. In contrast, the ResNet and MLP models showed relatively flat curves, indicating limited improvement. Nonetheless, the fact that models with lower training loss tended to stop earlier suggests that this decrease in error may not have fully transferred to the validation data.

**Figure 6.**
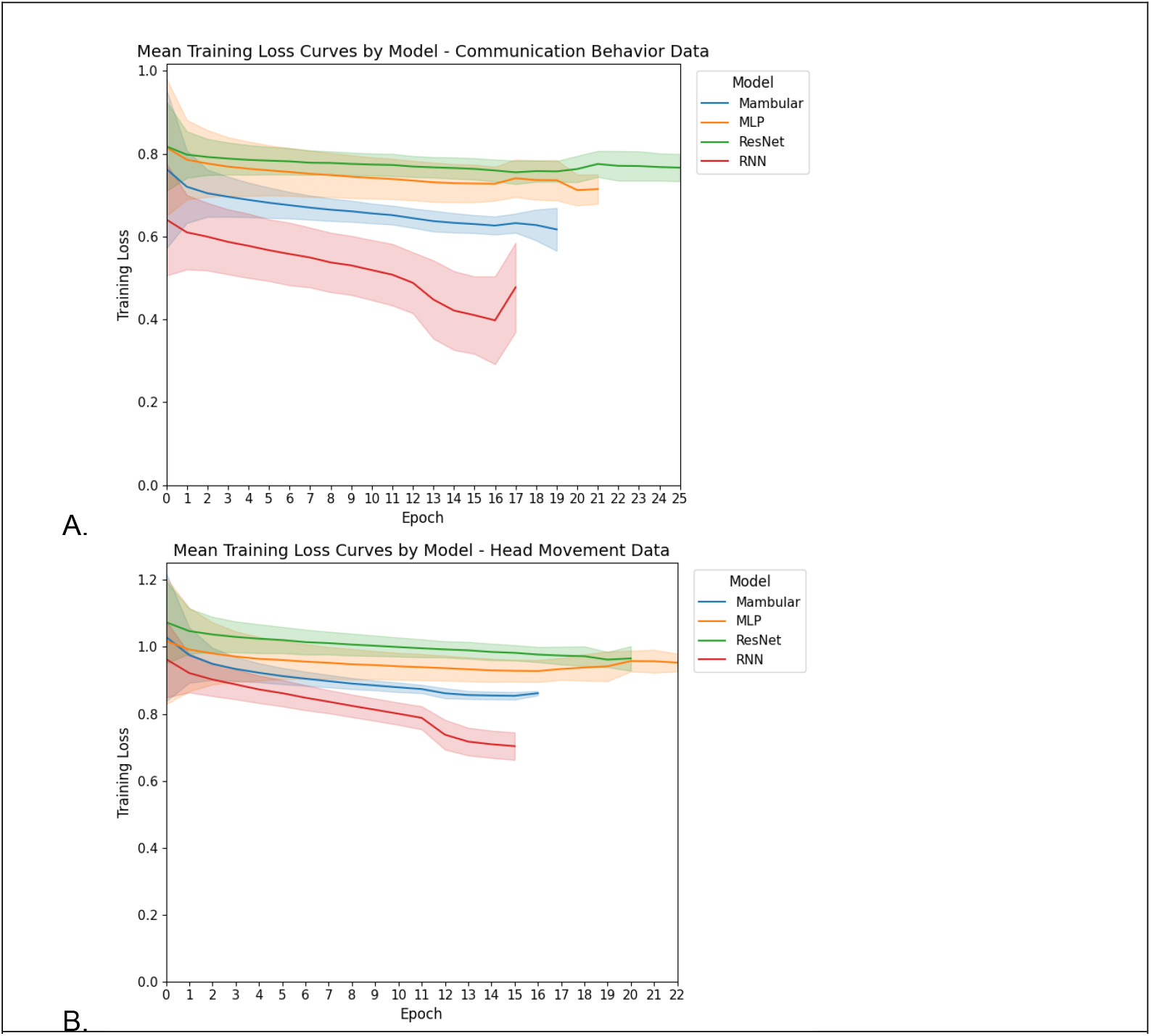
Training curves for four deep learning models on (A) communication behavior and (B) head movement data. Each epoch represents one complete pass through the dataset. Decreasing levels of training loss correspond to improved model performance. Training for each model ended once performance stopped improving on unseen data due to the “early stopping” option.

Figure 6B presents the mean training loss curves for the head movement data. As with the communication behavior data, the RNN and Mambular models showed the largest decreases in error, although the effect was less pronounced. These models also tended to stop earlier, and all models started and ended with higher loss values than in the communication behavior data, suggesting comparatively weaker performance.

In Figure 7A, the macro-F1 scores are plotted for both classical machine learning and deep learning models when applied to the communication behavior data, alongside dummy models shown in blue. The deep learning models consistently performed above the dummy baseline, often achieving scores greater than 0.6. The classical models also exceeded the dummy, but their scores were generally lower, more frequently in the 0.4-0.6 range. Importantly, these results varied across subjects: subject 1 showed uniformly lower scores across all model types, while subject 3 was the only case where some classical models did not surpass the dummy baseline. Among the deep learning models, the RNN achieved the strongest performance, ranking highest in 8 of the 9 subjects. For the classical models, the linear SVM performed best, outperforming the others in 7 out of 9 subjects.

**Figure 7.**
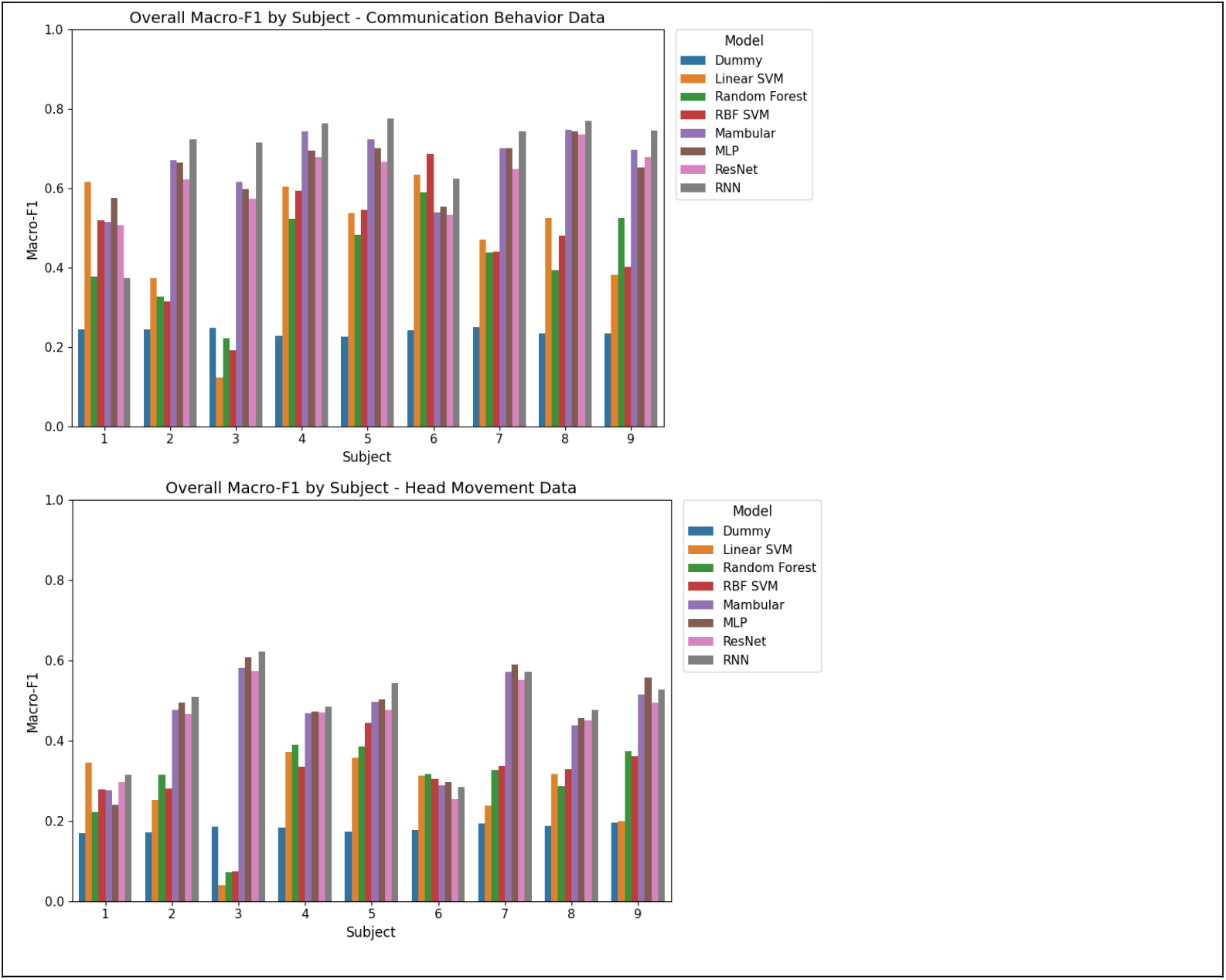
Macro-F1 scores across 9 test subjects for (A) communication behavior and (B) head movement data. Each model was trained on 8 subjects and tested using the remaining one. Colored bars represent different models, with blue indicating the dummy baseline. Linear SVM, Random Forest, and RBF SVM are classical machine learning models, while Mambular, MLP, ResNet, and RNN are deep learning models.

Figure 7B shows the macro-F1 scores for head movement data across classical and deep learning models, alongside the dummy baseline. As with the communication behavior data, all models outperformed the dummy model, and the deep learning models outperformed the classical machine learning. Overall, scores were about 0.1 to 0.2 points lower than in the communication behavior data. This difference is partly attributable to the increased difficulty of the head movement task for the model, which involved five classes instead of four, as reflected in the lower dummy scores. Note that the two subjects who performed poorly on the communication behavior data also had lower scores for the head movement behavior. Altogether, the deep learning models performed at a similar level in this task, with the RNN leading in 6 out of 9 subjects. In contrast, neither the linear SVM model nor any of the other classical machine learning models consistently outperformed the rest.

Tables 3 and 4 present the Macro-F1, Weighted-F1, and class-level F1 scores for each model type. Overall trends in Macro-F1 are consistent across tasks. The dummy model showed the weakest performance across all communication behaviors and nearly all head movement behaviors (the lower bound of the Linear SVM’s confidence interval only matched the dummy’s upper bound in the latter case). Overlap between classical and deep learning models was minimal, with a maximum of 0.02, supporting the conclusion that deep learning models outperformed classical machine learning models when all classes are considered equally. Within categories (classical vs. classical; deep learning vs. deep learning), performance was more comparable. RNNs achieved the highest mean scores for both communication and head movement, and Linear SVM led among communication behaviors. Nonetheless, these differences were small and fell within the overlapping intervals of the other models in the same category. Weighted-F1 scores showed the same trends but had higher means. With hyperparameter tuning, these means could further improve; in addition, differences within categories might become evident.

**Table 3:** Communication Behavior F1 Scores with 95% Confidence Intervals.

|  |  |  |  |  |  |  |
| --- | --- | --- | --- | --- | --- | --- |
| Communication Behavior Model | F1-Macro Score | F1-Weighted Score | Listen F1 Score | Talk F1 Score | Watch Video F1 Score | Other F1 Score |
| Dummy | 0.24 +/- 0.01 | 0.26 +/- 0.01 | 0.28 +/- 0.02 | 0.20 +/- 0.03 | 0.19 +/- 0.01 | 0.27 +/- 0.01 |
| Linear SVM | 0.47 +/- 0.12 | 0.51 +/- 0.13 | 0.57 +/- 0.17 | 0.34 +/- 0.12 | 0.39 +/- 0.21 | 0.60 +/- 0.14 |
| Random Forest | 0.43 +/- 0.09 | 0.47 +/- 0.09 | 0.48 +/- 0.11 | 0.38 +/- 0.06 | 0.27 +/- 0.21 | 0.59 +/- 0.11 |
| RBF SVM | 0.46 +/- 0.11 | 0.50 +/- 0.11 | 0.55 +/- 0.16 | 0.38 +/- 0.09 | 0.37 +/- 0.21 | 0.56 +/- 0.13 |
| Mambular | 0.66 +/- 0.07 | 0.68 +/- 0.07 | 0.69 +/- 0.08 | 0.52 +/- 0.09 | 0.72 +/- 0.12 | 0.72 +/- 0.08 |
| MLP | 0.65 +/- 0.05 | 0.67 +/- 0.05 | 0.67 +/- 0.07 | 0.53 +/- 0.10 | 0.71 +/- 0.07 | 0.71 +/- 0.08 |
| ResNet | 0.63 +/- 0.06 | 0.65 +/- 0.06 | 0.66 +/- 0.07 | 0.51 +/- 0.09 | 0.66 +/- 0.09 | 0.68 +/- 0.08 |
| RNN | 0.69 +/- 0.10 | 0.71 +/- 0.09 | 0.71 +/- 0.11 | 0.57 +/- 0.08 | 0.75 +/- 0.21 | 0.75 +/- 0.08 |

**Table 4:** Head Movement F1 Scores with 95% Confidence Intervals.

| Table 4: Head Movement F1 Scores with 95% Confidence Intervals |  |  |  |  |  |  |  |
| --- | --- | --- | --- | --- | --- | --- | --- |
| Head Movement Model | F1-Macro Score | F1-Weighted Score | Face Down F1 Score | Face Forward F1 Score | Turn Left F1 Score | Turn Right F1 Score | Other F1 Score |
| Dummy | 0.18 +/- 0.01 | 0.21 +/- 0.01 | 0.19 +/- 0.03 | 0.11 +/- 0.06 | 0.21 +/- 0.04 | 0.21 +/- 0.03 | 0.19 +/- 0.02 |
| Linear SVM | 0.27 +/- 0.08 | 0.34 +/- 0.12 | 0.61 +/- 0.20 | 0.00 +/- 0.00 | 0.21 +/- 0.11 | 0.31 +/- 0.17 | 0.21 +/- 0.15 |
| Random Forest | 0.30 +/- 0.08 | 0.36 +/- 0.10 | 0.60 +/- 0.19 | 0.12 +/- 0.15 | 0.23 +/- 0.13 | 0.28 +/- 0.10 | 0.27 +/- 0.10 |
| RBF SVM | 0.31 +/- 0.08 | 0.37 +/- 0.10 | 0.61 +/- 0.22 | 0.13 +/- 0.20 | 0.20 +/- 0.12 | 0.28 +/- 0.13 | 0.30 +/- 0.15 |
| Mambular | 0.46 +/- 0.08 | 0.54 +/- 0.06 | 0.84 +/- 0.08 | 0.21 +/- 0.25 | 0.47 +/- 0.18 | 0.46 +/- 0.11 | 0.31 +/- 0.11 |
| MLP | 0.47 +/- 0.10 | 0.55 +/- 0.07 | 0.84 +/- 0.11 | 0.23 +/- 0.26 | 0.48 +/- 0.20 | 0.49 +/- 0.09 | 0.31 +/- 0.11 |
| ResNet | 0.45 +/- 0.08 | 0.53 +/- 0.06 | 0.81 +/- 0.08 | 0.22 +/- 0.25 | 0.46 +/- 0.18 | 0.44 +/- 0.11 | 0.32 +/- 0.11 |
| RNN | 0.48 +/- 0.09 | 0.57 +/- 0.06 | 0.84 +/- 0.07 | 0.23 +/- 0.25 | 0.50 +/- 0.18 | 0.47 +/- 0.11 | 0.37 +/- 0.14 |

Note: Confidence intervals for both tables were calculated using the t-distribution (df = 8), appropriate for the small sample size (n = 9 subjects).

Breaking results down by condition highlighted variation in class-level performance. Talking was consistently harder to classify than other communication behaviors, while “face down” yielded the highest F1 score among head movement conditions and “face forward” the lowest. Once again, there was notable overlap within categories of models, but not across categories, further underscoring the superior performance of deep learning models. This finding is consistent with their ability to distinguish more complicated patterns in large datasets.

## Conclusion

From an experimental perspective, natural conversation between multiple participants represents a very uncontrolled environment. As a result, prior to any determination of listener intent based on head movements, a consistent framework of labeled behavioral annotations must be completed. The results presented here demonstrate that 1) multiple annotators can consistently identify head movement and communication behaviors across multiple hours of participant behavior, and 2) head movements captured by hearing-aid based accelerometers can be used to accurately classify head movement patterns distinct to different communication patterns.

Both classic machine and deep learning models were able to successfully distinguish between communication and head-movement behaviors, as demonstrated by their consistent ability to outperform a dummy baseline. Although mean performance differences within categories were modest, deep learning approaches outperformed classical machine learning methods. This outcome highlights their advantage in capturing more complex patterns within large datasets. With further hyperparameter tuning, overall accuracy and sensitivity to subtle differences across deep learning architectures are likely to improve.

This work also establishes a foundation for developing models that integrate sequential information. In other words, they would make predictions based not only on current inputs but also on temporal context from both past ones. Additionally, combining the communication behavior and head movement models into a single framework offers a promising avenue for enhancing predictive power. Together, these directions can contribute to more accurate detection of listener intent in real-world contexts and support the design of increasingly effective hearing devices.

## References

Appleton, J. (2022). What is important to your hearing aid clients… and are they satisfied? Hearing Review 29, 10–16.

Best, V., Roverud, E., Streeter, T., Mason, C. R., and Kidd, G. (2017). The Benefit of a Visually Guided Beamformer in a Dynamic Speech Task. Trends Hear 21, 2331216517722304. doi: 10.1177/2331216517722304

Breiman, L. (2001). Random Forests. Machine Learning 45, 5–32. doi: 10.1023/A:1010933404324

Cortes, C., and Vapnik, V. (1995). Support-vector networks. Machine Learning 20, 273–297. doi: 10.1007/BF00994018

Desai, N., Beukes, E. W., Manchaiah, V., Mahomed-Asmail, F., and Swanepoel, D. W. (2024). Consumer Perspectives on Improving Hearing Aids: A Qualitative Study. Am J Audiol 33, 728–739. doi: 10.1044/2024_AJA-23-00245

Doclo, S., Gannot, S., Moonen, M., and Spriet, A. (2010). Acoustic Beamforming for Hearing Aid Applications. Available at: https://api.semanticscholar.org/CorpusID:56623464

Elman, J. L. (1990). Finding structure in time. Cognitive Science 14, 179–211. doi: 10.1016/0364-0213(90)90002-E

Favre-Felix, A., Graversen, C., Dau, T., and Lunner, T. (2017). Real-time estimation of eye gaze by in-ear electrodes. Annu Int Conf IEEE Eng Med Biol Soc 2017, 4086–4089. doi: 10.1109/EMBC.2017.8037754

Favre-Félix, A., Graversen, C., Hietkamp, R. K., Dau, T., and Lunner, T. (2018). Improving Speech Intelligibility by Hearing Aid Eye-Gaze Steering: Conditions With Head Fixated in a Multitalker Environment. Trends in Hearing 22, 1–13. doi: 10.1177/2331216518814388

He, K., Zhang, X., Ren, S., and Sun, J. (2016). Deep Residual Learning for Image Recognition., in 2016 IEEE Conference on Computer Vision and Pattern Recognition (CVPR), 770–778. doi: 10.1109/CVPR.2016.90

Hladek, L., and Seeber, B. U. (2019). Behavior and Speech Intelligibility in a Changing Multi-talker Environment., 7640–7645.

Hochreiter, S., and Schmidhuber, J. (1997). Long Short-Term Memory. Neural Comput 9, 1735–1780. doi: 10.1162/neco.1997.9.8.1735

Ibrahim, I., Parsa, V., Macpherson, E., and Cheesman, M. (2013). Evaluation of Speech Intelligibility and Sound Localization Abilities with Hearing Aids Using Binaural Wireless Technology. Audiol Res 3, e1. doi: 10.4081/audiores.2013.e1

J. G. Desloge, W. M. Rabinowitz, and P. M. Zurek (1997). Microphone-array hearing aids with binaural output.I. Fixed-processing systems. IEEE Transactions on Speech and Audio Processing 5, 529–542. doi: 10.1109/89.641298

Jeni, L. A., Cohn, J. F., and Kanade, T. (2017). Dense 3D Face Alignment from 2D Video for Real-Time Use. Image Vis Comput 58, 13–24. doi: 10.1016/j.imavis.2016.05.009

Kidd, G. (2017). Enhancing Auditory Selective Attention Using a Visually Guided Hearing Aid. J Speech Lang Hear Res 60, 3027–3038. doi: 10.1044/2017_JSLHR-H-17-0071

Kidd, G., Favrot, S., Desloge, J. G., Streeter, T. M., and Mason, C. R. (2013). Design and preliminary testing of a visually guided hearing aid. J Acoust Soc Am 133, EL202–207. doi: 10.1121/1.4791710

Kidd, G., Mason, C. R., Best, V., and Swaminathan, J. (2015). Benefits of Acoustic Beamforming for Solving the Cocktail Party Problem. Trends Hear 19, 2331216515593385. doi: 10.1177/2331216515593385

Kohavi, R. (1995). A Study of Cross-Validation and Bootstrap for Accuracy Estimation and Model Selection., in International Joint Conference on Artificial Intelligence. Available at: https://api.semanticscholar.org/CorpusID:2702042

Lausberg, H., and Sloetjes, H. (2009). Coding gestural behavior with the NEUROGES--ELAN system. Behav Res Methods 41, 841–849. doi: 10.3758/BRM.41.3.841

Lu, H., McKinney, M. F., Zhang, T., and Oxenham, A. J. (2021). Investigating age, hearing loss, and background noise effects on speaker-targeted head and eye movements in three-way conversations. J Acoust Soc Am 149, 1889. doi: 10.1121/10.0003707

Nasreddine, Z. S., Phillips, N. A., Bédirian, V., Charbonneau, S., Whitehead, V., Collin, I., et al. (2005). The Montreal Cognitive Assessment, MoCA: a brief screening tool for mild cognitive impairment. J Am Geriatr Soc 53, 695–699. doi: 10.1111/j.1532-5415.2005.53221.x

Neher, T., Wagener, K. C., and Latzel, M. (2017). Speech reception with different bilateral directional processing schemes: Influence of binaural hearing, audiometric asymmetry, and acoustic scenario. Hear Res 353, 36–48. doi: 10.1016/j.heares.2017.07.014

Rosenblatt, F. (1958). The perceptron: A probabilistic model for information storage and organization in the brain. Psychological Review 65, 386–408. doi: 10.1037/h0042519

Roverud, E., Best, V., Mason, C. R., Streeter, T., and Kidd, G. (2018). Evaluating the Performance of a Visually Guided Hearing Aid Using a Dynamic Auditory-Visual Word Congruence Task. Ear Hear 39, 756–769. doi: 10.1097/AUD.0000000000000532

Russell, S. J., Russell, S., and Norvig, P. (2020). Artificial Intelligence: A Modern Approach. Pearson. Available at: https://books.google.com/books?id=koFptAEACAAJ

Smeds, K., Wolters, F., and Rung, M. (2015). Estimation of Signal-to-Noise Ratios in Realistic Sound Scenarios. J Am Acad Audiol 26, 183–196. doi: 10.3766/jaaa.26.2.7

Thielmann, A. F., Kumar, M., Weisser, C., Reuter, A., Säfken, B., and Samiee, S. (2024). Mambular: A Sequential Model for Tabular Deep Learning. Available at: https://openreview.net/forum?id=wElgE9qBb5 (Accessed September 26, 2025).

Van den Bogaert, T., Klasen, T. J., Moonen, M., Van Deun, L., and Wouters, J. (2006). Horizontal localization with bilateral hearing aids: without is better than with. J Acoust Soc Am 119, 515–526. doi: 10.1121/1.2139653

Wagener, K. C., Hansen, M., and Ludvigsen, C. (2008). Recording and classification of the acoustic environment of hearing aid users. J Am Acad Audiol 19, 348–370. doi: 10.3766/jaaa.19.4.7

Wolters, F., Smeds, K., Schmidt, E., Christensen, E. K., and Norup, C. (2016). Common Sound Scenarios: A Context-Driven Categorization of Everyday Sound Environments for Application in Hearing-Device Research. J Am Acad Audiol 27, 527–540. doi: 10.3766/jaaa.15105

Wu, Y.-H., and Bentler, R. A. (2012). Do older adults have social lifestyles that place fewer demands on hearing? J Am Acad Audiol 23, 697–711. doi: 10.3766/jaaa.23.9.4

